# IFNγ-Driven Inflammatory Responses in the Nasal Mucosa Drive Influenza Virus Shedding and Transmission

**DOI:** 10.64898/2026.01.06.698034

**Authors:** Stacey Bartlett, Hedy L. Rocha, Leon L. Hsieh, Stephen T. Yeung, Chan Wang, Grace O. Ciabattoni, Cynthia A. Loomis, Huilin Li, Benjamin tenOever, Mila B. Ortigoza

**Affiliations:** Department of Medicine, NYU Grossman School of Medicine, New York, NY, 10016, USA; Department of Microbiology, NYU Grossman School of Medicine, New York, NY, 10016, USA; Division of Biostatistics, Department of Population Health, NYU Grossman School of Medicine, New York, 10016, NY, USA; Department of Pathology, NYU Grossman School of Medicine, New York, New York, USA; Department of Medicine, Division of Infectious Disease, Weill Cornell Medicine, New York, NY, 10021, USA

**Author notes:** Corresponding author Mila B. Ortigoza, MD Department of Medicine, NYU Grossman School of Medicine Alexandria Center for Life Science, West Tower 430 East, 29th Street, Suite 308 New York, NY 10016.

## Abstract

What determines host infectiousness during influenza A virus (IAV) infection remains a fundamental unanswered question in virology. While upper respiratory tract (URT) replication is necessary for transmission, it’s insufficient to explain host-to-host variation in contagiousness. Using the infant mouse model of influenza transmission, we show that viruses containing H3 hemagglutinin shed at higher levels than non-H3-containing viruses, despite comparable URT replication. H3-containing virus infection was associated with higher URT inflammation, characterized by increased cytokine production, immune cell recruitment, and mucus hypersecretion, which directly correlated with shedding efficiency. Transcriptomic profiling identified enrichment of interferon-stimulated gene programs, with a dominant interferon gamma (IFNγ) signature during H3 infections. Functional studies demonstrated that IFNγ deficiency reduced mucus production, shedding, and transmission, whereas IFNγ supplementation restored these phenotypes. Together, these findings identify IFNγ-driven mucosal inflammation key host determinant of influenza infectiousness, reframing contagiousness as a consequence of host inflammatory clearance rather than viral replication alone.

## Introduction

Influenza A virus (IAV) transmission is a complex process shaped by the interplay between viral characteristics and host physiological responses to infection. Efficient spread requires infected hosts (index) to shed sufficient amounts of infectious virus from the upper respiratory tract (URT) under conditions that enable efficient exposure to susceptible hosts (contacts). Although viral replication is essential for this process, it alone cannot fully account for the marked variability in shedding efficiency and transmission observed across viral strains and individuals. Epidemiological studies in humans show that seasonal H3N2 viruses generally cause higher infection attack rates, a proxy for transmissibility, than H1N1 viruses^1^. Moreover, transmission is highly heterogeneous, with a small fraction of individuals (∼20%) responsible for the majority (∼80%) of secondary transmission events, a phenomenon known as “superspreading”^2–4^. Together, these observations highlight the importance of host-specific factors in shaping contagiousness and raise a key unresolved question: ***what makes a host infectious?***

Human challenge studies have shown that IAV can be transmitted through multiple routes, including respiratory droplets (>5µm), aerosols (<5µm), and direct contact with respiratory secretions. Across these routes, host infectiousness is strongly influenced by the magnitude, timing, and duration of viral shedding, which vary widely among individuals and viral strains^4^. Although viral load in the URT is a prerequisite for efficient shedding, it is insufficient to fully explain differences in shedding efficiency or transmission potential. Increasing evidence points to local host factors, particularly the immune response and the URT mucosal environment, as key regulators of virus shedding efficiency^5–7^.

Infection of the URT mucosa rapidly triggers a localized inflammatory response that is central to antiviral defense. This response includes epithelial activation, cytokine and chemokine secretion, recruitment of immune cells, and stimulation of goblet cell hyperplasia and mucin gene expression, resulting in increased mucus production^8^. Interferons (IFNs) are critical orchestrators of this process. Beyond restricting viral replication, IFNs modulate epithelial function, amplify inflammatory signaling, and promote mucus secretion^9,10^. Together, these changes enhance mucociliary flow, a host-protective mechanism that traps virus within mucus and expels it from the airway, thereby limiting progression of infection to the lower respiratory tract (LRT).

However, these same expulsive processes that protect the host may also increase opportunities for virus transmission. Enhanced mucus production and accelerated mucociliary clearance facilitate the release of virus-laden secretions into the environment, increasing the likelihood of exposure to nearby contacts. Consistent with this concept, studies in the ferret model have linked more robust early innate immune responses with higher peak viral shedding, increased nasal discharge, and more efficient transmission^5^. Similarly, human studies show that nasal inflammation and congestive symptoms correlate with increased mucus production, virus shedding, and transmission^6^. Collectively, these findings suggest that URT inflammation promotes viral clearance through shedding while simultaneously increasing transmission potential as an unintended consequence of an effective local immune response.

Mechanistic insight into how specific inflammatory pathways modulate transmission has been limited in part by constraints of available animal models. Ferrets and guinea pigs recapitulate many features of human influenza infection and transmission, but their use is restricted by cost, housing requirements, and a limited availability of species-specific reagents^11^. Mice, in contrast, are cost-effective and supported by extensive genetic and immunological tools, yet adult mice infected with IAV rarely transmit efficiently to cohoused contacts^12^. To overcome these limitations, we established an infant mouse model that supports robust, reproducible IAV transmission under close-contact conditions, resembling transmission dynamics observed in children within household settings^7,12^. Infant mice exhibit high baseline expression of both α2,3- and α2,6-linked sialic acid receptors in the URT epithelium, critical for IAV entry, whereas adults display lower sialic acid expression^12^. These features, along with the augmented susceptibility of the infant respiratory tract to pathogens in general, make this model uniquely powerful to dissect the interplay between viral factors, host inflammatory responses, and transmission dynamics.

In this study, we use the infant mouse model to define how host inflammatory responses modulate IAV shedding and transmission. By comparing viruses that differ in transmissibility, we show that H3-containing viruses shed at higher levels and transmit more efficiently than non-H3 viruses, despite comparable replication in the URT. H3 infection was consistently associated with heightened URT inflammation, characterized by increased cytokine production, immune cell recruitment, and pronounced mucus hypersecretion, which closely tracked with shedding magnitude. Because inflammation expands the mucus layer and enhances mucociliary flow, this hypersecretory state promotes viral expulsion from the URT and increases opportunities for transmission. Integrating transcriptomic and functional approaches, we identify interferon gamma (IFNγ) as a central regulator of this process. IFNγ deficiency reduced mucus production, viral shedding, and transmission, whereas IFNγ supplementation restored these outcomes without increasing URT viral loads. We propose that IFNγ-driven mucus rich inflammation in the URT is not merely permissive but actively drives infectiousness, reframing influenza transmission as a host-regulated consequence of antiviral clearance rather than a direct function of viral replication. Collectively, these findings establish IFNγ-dependent mucosal inflammation as key regulators of IAV transmission. These insights suggest new opportunities to identify hosts at increased risk of superspreading and to explore modulation of mucosal inflammatory responses as a means of limiting viral transmission.

## Results

### Influenza viruses shed and transmit at different efficiencies

To define determinants of IAV transmissibility, we employed the infant mouse model to systematically compare URT shedding and transmission efficiency across representative IAV strains. C57BL/6 infant mice (4-7 days old) were infected intranasally with 300 plaque forming units (PFU) using a low-volume inoculum to confine infection to the URT^12–15^. Following infection, index pups were returned to the cage with the dam and naïve littermates (contacts). Viral shedding from index pups was monitored daily by gently dipping the nares of each pup into viral medium to collect loose virus released at the URT surface. Transmission was assessed in contact pups by retrograde URT lavage at sacrifice at 4 days post-contact, a timepoint at which >90% transmission is typically achieved for transmissible viruses in this model^7^. Infectious titers from daily shedding samples and terminal URT lavages were quantified by standard plaque assay (**Fig.1A**).

**Figure 1:**
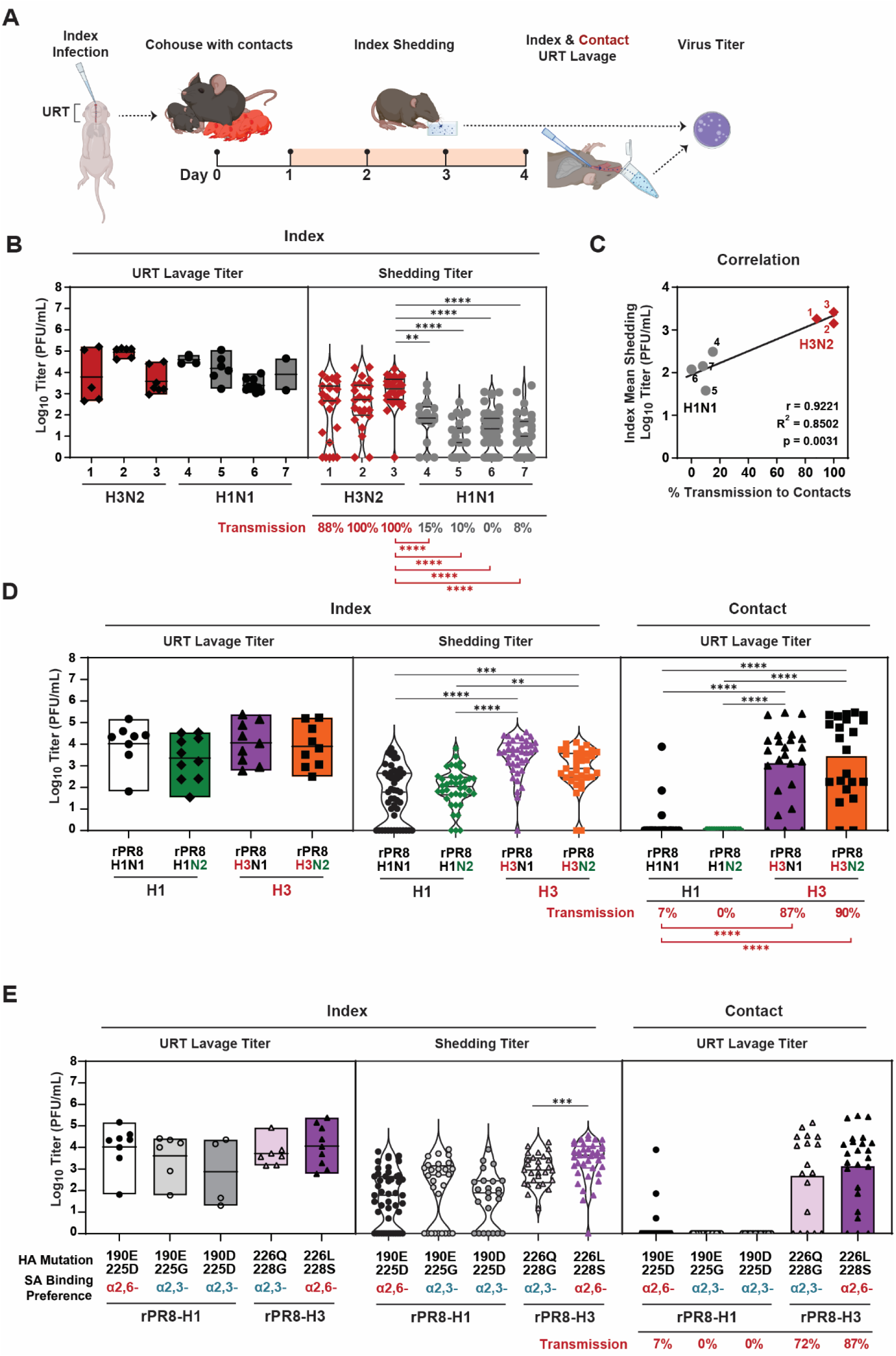
Enhanced viral shedding of H3-containing viruses in mice. (A) <u>Schematic of the infant mouse transmission model</u>. C57BL/6J index pups were intranasally infected at day 0 with 300 PFU of the indicated influenza virus strains and immediately cohoused with uninfected littermates (contacts) for 4 days. Daily shed secretions (days 1–4) were collected from index pups and quantified by plaque assay. On day 4, both index and contact pups were sacrificed, and URT lavages were collected to assess infectious viral titers by plaque assay. (B) <u>Shedding and transmission of diverse influenza virus strains</u>. Infectious titers from day 4 URT lavages (left) and cumulative shedding from days 1–4 (right) were quantified by plaque assay. Each point represents an individual mouse; median values are indicated. Transmission outcomes for contact pups are shown in red. Virus strain identities are listed in Table 1: (1) Pan/99, (2) HK/68, (3) X-31, (4) PR/8, (5) WSN, (6) Cal/09, (7) Bris/07. (C) <u>Shedding magnitude strongly correlates with transmission efficiency</u>. Mean shedding titers from index pups were plotted against transmission rates to contacts for each virus strain. Pearson’s correlation (*r*), best fit linear regression, and goodness of fit (*R*^2^) were generated in GraphPad Prism 10. (D) <u>Recombinant viruses reveal HA-specific contributions to shedding and transmission</u>. Viral titers from day 4 URT lavage (left), cumulative shedding from days 1–4 (middle), and day 4 URT titers from contact pups (right) were quantified. Each point represents an individual mouse; median values are shown. Transmission frequencies are indicated in red. (E) <u>Altered sialic acid–binding preference modestly affects shedding and transmission</u>. Day 4 URT lavage titers and cumulative shedding (days 1–4) were quantified for recombinant viruses engineered to favor α2,3- or α2,6-linked sialic acid binding. Transmission outcomes to contact pups are indicated in red. <u>Statistics</u>. Differences among group medians were analyzed by Kruskal–Wallis test. Experiments represent ≥2 independent biological replicates. Significance is denoted as *P < 0.05, **P < 0.01, ***P < 0.001, ****P < 0.0001. Cartoons created with BioRender.com; graphs generated using GraphPad Prism 10.

Among the panel of viruses tested, H3N2 strains consistently shed at significantly higher titers than H1N1 strains, even though both subtypes achieved equivalent URT viral loads by the end of the experiment, as measured by nasal lavage^7,16^. This indicates that viral subtype can influence shedding efficiency independently of URT replication dynamics (**Fig.1B and Sup. Table 1**). In direct contact transmission experiments, when indexes are cohoused with contacts for 4 days, H3N2-infected index pups transmitted virus to a greater proportion of contacts compared to H1N1-infected pups. Importantly, the magnitude of viral shedding in index pups strongly correlated with transmission success to contacts, suggesting that shedding titers are a key predictor of contagiousness (**Fig.1C**).

These findings are consistent with epidemiological patterns observed in humans. H3N2 viruses have frequently predominated in seasonal influenza outbreaks over recent decades and have been the predominant circulating subtype in 62% of the past 29 influenza seasons, a pattern often associated with higher transmission and disease burden compared with H1N1 viruses^1,17–19^ (**Sup. Fig 1A)**. Together, these observations support our experimental findings and suggest that H3N2 viruses possess an inherent advantage in shedding and transmission potential.

To identify viral determinants underlying these differences in shedding and transmission, we focused on two well-characterized model strains: A/Puerto Rico/8/1934 (H1N1; A/PR/8) and its reassortant derivative A/X-31 (H3N2). These viruses share six internal gene segments but differ in their surface glycoproteins, hemagglutinin (HA) and neuraminidase (NA), with A/X-31 incorporating the HA and NA from the 1968 pandemic strain A/Hong Kong/1/1968 (H3N2, A/HK/68). This defined genetic relationship enabled isolation of the specific contributions of HA and NA to viral shedding and transmission. Using reverse genetics, we generated a panel of recombinant viruses containing A/PR/8 internal genes and H3 and/or N2 from A/HK/68 (rPR8, rPR8-N2, rPR8-H3, rPR8-H3N2)^20,21^ (**Sup. Fig 1B**). Viruses were rescued in 293T cells and propagated in embryonated chicken eggs. All recombinant viruses exhibited comparable infectivity and robust replication in the URT (**Sup. Fig 1C and Sup. Table 2**). Despite similar URT viral loads at 4dpi, pups infected with rPR8-H3N2 and rPR8-H3 shed at significantly higher titers than those infected with rPR8 and rPR8-N2 across days 1-4 post infection, resulting in markedly higher transmission efficiency to contacts (**Fig.1D**). These results implicate H3 as a key determinant of shedding efficiency from the URT. Notably, shedding by rPR8-H3N2 and rPR8-H3 peaked within the first 2 days post infection (dpi), a period critical for transmission (**Sup. Fig. 1D**). Reducing the inoculum of rPR8-H3 tenfold significantly impaired shedding and transmission (**Sup. Fig. 1E**), demonstrating that sufficient URT replication is necessary but not alone sufficient to drive efficient spread. Conversely, extending index-contact exposure from 4 to 7 days increased transmission of rPR8 (7% to 49%) and rPR8-N2 (0% to 44%) without increasing cumulative shedding titers (**Sup. Fig. 1F**), indicating that prolonged exposure can partially compensate for inefficient shedding. Together, these findings indicate that reduced transmission of H1N1 viruses in this model reflects the inherent complexity of transmission dynamics, where successful spread is driven by the magnitude and timing of index shedding and the duration of contact exposure, rather than by viral replication alone.

Given the prominent contribution of H3 to enhanced shedding and transmission, we hypothesized whether differences in HA receptor binding preference could account for this phenotype. To assess receptor-specific contributions, we focused on the two major sialic acid (SA) linkages, α2,3- and α2,6-linked SA, that mediate IAV entry through HA engagement. Known amino acid substitutions, including E190D and G225D in H1 or Q226L and G228S in H3, are known to shift receptor binding preference from α2,3-SA (‘avian-like’) to α2,6-SA (‘human-like’)^22^. Because infant mice express both α2,3-SA and α2,6-SA in the URT^12^, this model provides an opportunity to directly evaluate the impact of SA preference on viral shedding and transmission. While α2,6-SA binding conferred a modest increase in shedding during rPR8-H3 infection, differences in transmission efficiency between α2,3- and α2,6-SA-binding viruses were not significant (**Fig.1E**). Notably, rPR8 exhibited reduced shedding and transmission efficiency regardless of SA binding preference, indicating that receptor specificity alone is insufficient to explain the enhanced shedding and transmission associated with H3-containing viruses. These findings suggest that additional HA-associated properties beyond receptor binding contribute to the increased transmission potential of H3N2 viruses.

### Transmissible viruses are more inflammatory in the URT

Building on our observation that highly transmissible viruses (HTVs), such as rPR8-H3N2 and rPR8-H3, shed and transmit more efficiently than low transmissible viruses (LTVs), such as rPR8 and rPR8-N2, we next sought to identify host factors contributing to these differences. Although viral determinants, particularly the presence of H3, clearly influence shedding capacity, our receptor-binding analyses indicated that HA-mediated entry alone was insufficient to explain the magnitude, timing, or efficiency of viral shedding and transmission. Moreover, substantial heterogeneity in transmission persists even among hosts infected with the same virus, with only a subset of individuals shedding and transmitting virus at high levels, highlighting a critical role for host physiology in shaping contagiousness^2,3^. Together, these results suggest that viral determinants such as HA modulate transmission through host pathways that remain incompletely defined. We therefore leveraged the infant mouse model to define how inflammatory responses within the URT contribute to viral shedding and transmission.

To dissect how immune activation influences transmissibility, we leveraged our isogenic panel of recombinant viruses. Infant mice were infected intranasally with either HTV (H3-containing viruses) or LTV strains (H1-containing viruses), and URT samples were collected at 2 dpi, corresponding to peak shedding. Nasal lavages were obtained to quantify secreted inflammatory mediators, and cellular fractions from URT were harvested for immune profiling (**Fig.2A**). Because all recombinant viruses replicated to comparable titers in the URT (**Fig.2B**), this approach enabled focused analysis of host responses associated with enhanced shedding and transmission.

**Figure 2:**
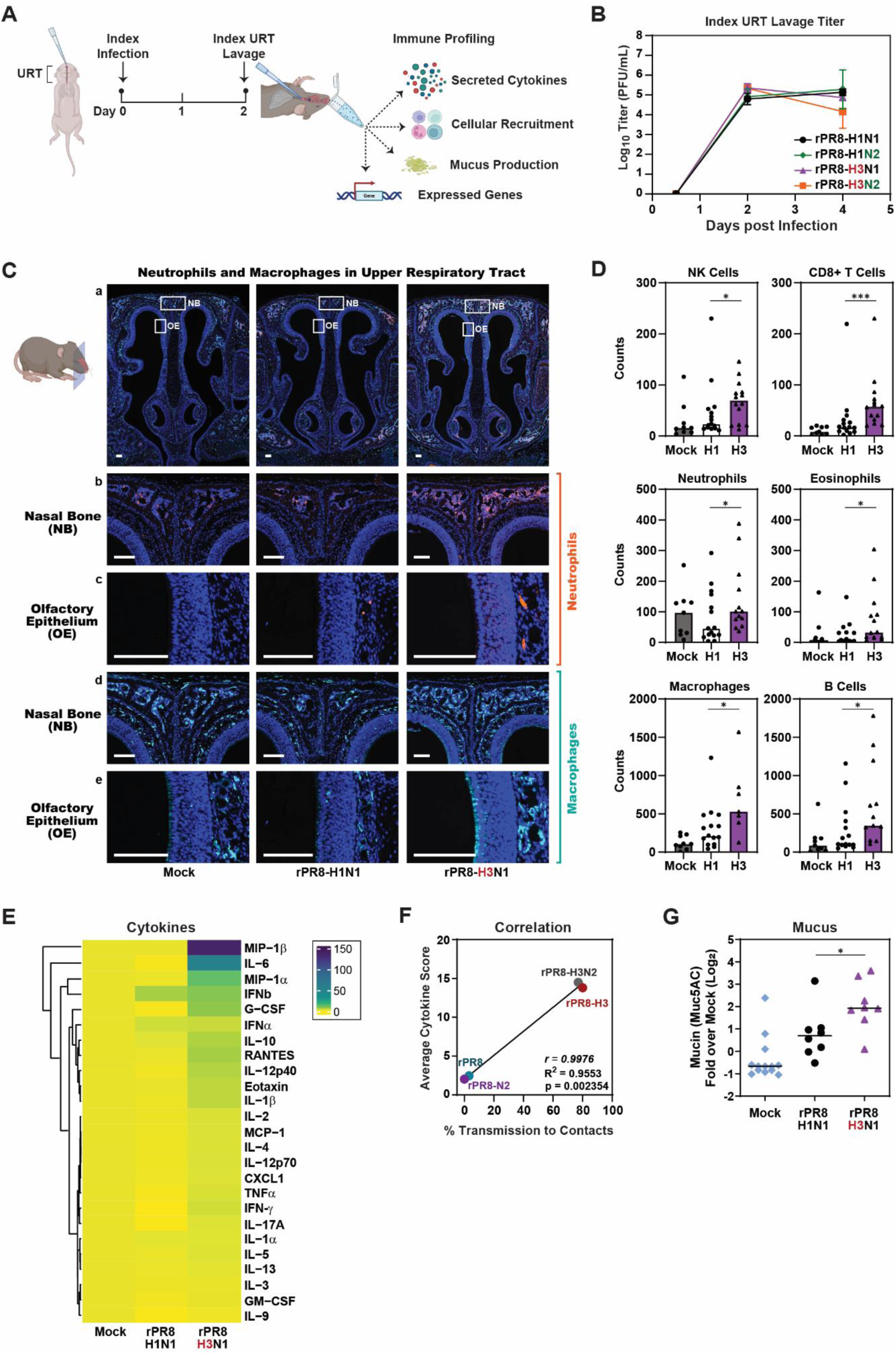
Inflammatory profiles of infant mice infected with highly transmissible H3-containing and low transmissible H1-containing influenza viruses. (A) <u>Schematic of immune profiling workflow</u>. Infant mice were intranasally infected with 300 PFU of the indicated influenza virus strains. At 2 dpi, pups were sacrificed and URT lavages were collected for comprehensive analysis, including viral titration by plaque assay, immune cell characterization by flow cytometry, cytokine quantification by Luminex, and mucus assessment by immunoblot. (B) <u>Viral titers in the URT</u>. Viral burden in URT lavage fluid at 2 and 4 dpi was quantified by plaque assay. Each point represents an individual mouse; error bars indicate SEM. (C) <u>Histological assessment of inflammatory cell recruitment</u>. Infant mice infected with rPR8-H3 or rPR8-H1 were sacrificed at 2 dpi. Heads were fixed in 4% paraformaldehyde, paraffin-embedded, sectioned through the nasopharynx, and stained for Ly6G (neutrophils) or F4/80 (macrophages). Scale bars = 100 μm. (D) <u>Flow cytometry of URT immune infiltrates</u>. URT lavages collected at 2 dpi were analyzed by flow cytometry. Cellular abundance of NK cells, CD8⁺ T cells, neutrophils, eosinophils, macrophages, and B cells are shown as the fraction of total live CD45⁺ cells. Median values are displayed. Gating was performed using FlowJo. (E) <u>Cytokine profiling of URT secretions</u>. Cytokines from URT lavages of rPR8-H3 or rPR8-H1 infected pups at 2 dpi were quantified using a Luminex multiplex assay. Values represent fold-change relative to mock. Heatmap was generated using RStudio. (F) <u>Correlation between cytokine responses and transmission efficiency</u>. Average cytokine levels from index pups infected with rPR8-H3 or rPR8-H1 were compared to transmission rates in contact pups. Pearson’s correlation (*r*), best fit linear regression, and goodness of fit (*R*^2^) were generated in GraphPad Prism 10. (G) <u>Mucus quantification in the URT</u>. URT lavages collected at 2 dpi were dotted on nitrocellulose and probed with anti-Muc5AC antibody. Signal intensity was normalized to positive control, and fold-change over mock is shown. Median values are indicated. <u>Statistics</u>. Differences between two groups were analyzed using the Mann–Whitney test. Differences among multi-group medians were analyzed by Kruskal–Wallis test. Experiments represent ≥2 independent biological replicates. Significance is denoted as *P < 0.05, **P < 0.01, ***P < 0.001, ****P < 0.0001. Cartoons created with BioRender.com; graphs generated using GraphPad Prism 10.

We first assessed whether H3-containing virus infection elicited stronger local inflammation than H1-containing virus infection. Given that immune cell infiltration provides a proximal readout of mucosal inflammatory activation, we characterized URT cellular populations using immunohistochemistry and flow cytometry. H3-infected mice exhibited significantly increased infiltration of neutrophils and macrophages compared with H1-infected mice (**Fig.2C-D**). Additional increases were observed in eosinophils, NK cells, B cells, and CD8+ T cells, whereas other immune subsets remained unchanged (**Sup. Fig. 2A-B)**, indicating that H3-containing infection induces a broader inflammatory response within the URT.

To determine whether immune cell recruitment was accompanied by corresponding changes in inflammatory signaling, we next quantified cytokines and chemokines in nasal lavages at 2 dpi. Both H3- and H1-infected mice mounted inflammatory responses relative to uninfected controls (**Fig.2E**). However, H3-containing infection was associated with significantly higher levels of multiple inflammatory mediators, including MIP-1α, MIP-1β, IL-6, G-CSF, CCL5, and IFNγ, compared to LTV infection (**Sup. Fig. 3A**). Many of these inflammatory mediators promote recruitment and activation of macrophages, neutrophils, and NK cells, consistent with the observed cellular influx. Accordingly, a composite cytokine score was significantly elevated during H3 infection relative to H1 infection, linking heightened inflammatory signaling with increased transmission capacity (**Fig.2F**).

**Figure 3:**
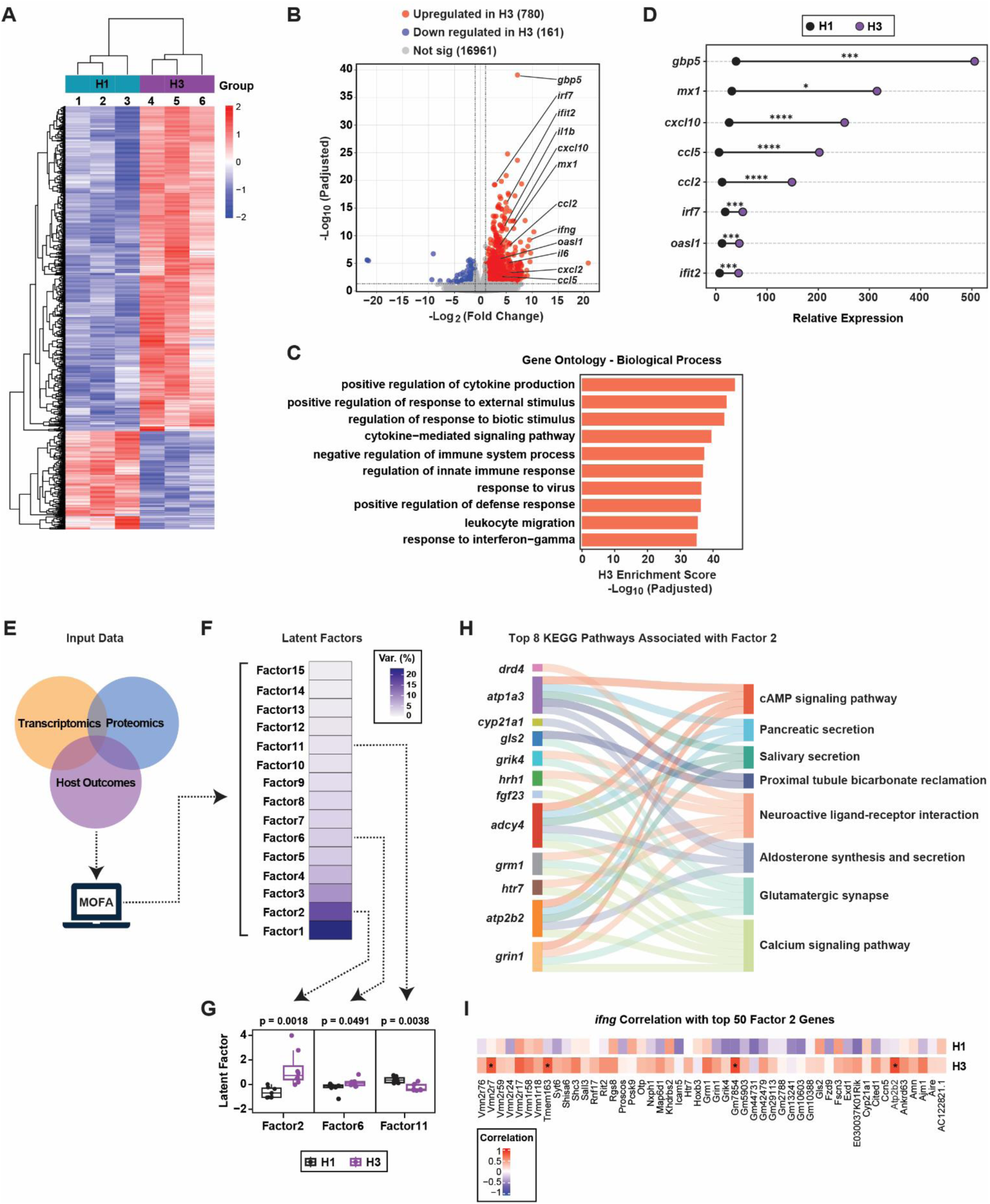
URT transcriptional programs induced by H3-containing viruses. Mice were intranasally infected with 300 PFU of rPR8-H3 or rPR8-H1. At 2 dpi, pups were sacrificed and URT lavages were collected. RNA was extracted from cellular fractions for bulk RNA sequencing. <u>Differential Gene Expression Analysis</u>. (**A**) Heatmap of differentially expressed genes (DEGs) between rPR8-H3 and rPR8-H1 infected mice. (**B**) Volcano plot depicting DEGs. A fold-change cutoff of ≥2 and an adjusted p-value of <0.05 were used to define significance. (**C**) Gene ontology (GO) enrichment analysis of the top 4,000 upregulated genes, highlighting pathways associated with inflammation, chemotaxis, and interferon signaling. (**D**) Independent qPCR validation of selected upregulated transcripts confirms enrichment of type I and type II interferon–responsive genes in rPR8-H3-infected mice. <u>Integration of Transcriptional, Cytokine, and Transmission Data</u>. (**E**) Schematic overview of the Multi-Omics Factor Analysis (MOFA) workflow integrating RNA-seq data, cytokine measurements, and transmission outcomes (transmissible vs. non-transmissible phenotypes) generated from the same URT samples. (**F**) MOFA identified 15 latent factors representing co-regulated biological programs that capture shared variation across samples. (**G**) Three factors (Factors 2, 6, and 11) significantly distinguished rPR8-H3 from rPR8-H1 infections, with Factor 2 showing the strongest and most consistent association with transmissibility. <u>Biological Interpretation of Factor 2</u>. (**H**) Sankey plot showing the top eight KEGG pathways enriched in Factor 2. These pathways converge on ion transport, ciliary regulation, and epithelial secretory function, driven by overlapping sets of IFNγ-associated genes. (**I**) Heatmap displaying the correlation coefficients between the top 50 genes with the highest positive loadings on Factor 2 and IFNγ abundance across samples, demonstrating strong alignment between rPR8-H3-induced transcriptional programs and IFNγ-driven inflammation. Graphs were generated using SRplot.

Given that mucosal inflammation is closely coupled to epithelial secretory responses, we next examined whether enhanced inflammation during H3-containing infection was associated with increased mucus production. Mucus plays a dual role during respiratory infection: entrapping viral particles while facilitating their clearance from the airway surface. Quantification of mucins in nasal lavages revealed that H3-containing infection induced significantly higher levels of the major secretory mucin Muc5AC compared with H1-containing infection (**Fig.2G**). Together, these findings suggest that enhanced transmissibility is associated with a hyperinflammatory URT environment during H3 infection, characterized by increased cytokine production, immune cell infiltration, and mucus secretion.

### Distinct inflammatory transcriptional programs characterize highly transmissible infections

To define the molecular pathways underlying the heightened inflammation and mucus secretion associated with HTV infection, we performed bulk RNA sequencing of URT lavages collected at 2 dpi from mice infected with H3- or H1-containing viruses. Differential gene expression and pathway enrichment analyses revealed distinct transcriptional programs associated with transmissibility, with H3-containing infections eliciting a strong induction of inflammatory, chemotactic, and interferon-stimulated gene (ISG) networks (**Fig.3A-C)**. Independent qPCR validation confirmed the RNA-seq findings and demonstrated enrichment of both type I and type II IFN responses during H3-containing infection. Notably, canonical IFNγ-responsive genes such as *gbp5*, *cxcl10*, and *ccl5* were among the most highly upregulated transcripts, consistent with enhanced recruitment of phagocytes, CD8+ T-cells, NK cells and amplification of local mucosal inflammation. In parallel, broadly antiviral ISGs including *mx1*, *ccl2*, *irf7*, *oasl1*, and *ifit1* were also elevated, reinforcing a robust antiviral state that may facilitate viral expulsion from the URT (**Fig.3D**). Together, these transcriptional signatures indicate that H3-containing infection induces a coordinated IFNγ-skewed inflammatory program associated with enhanced viral shedding.

To further resolve the biological programs linked to contagiousness, we integrated transcriptional profiles with cytokine abundance and transmission outcomes using Multi-Omics Factor Analysis (MOFA). This blinded approach summarizes transcriptomic variation into latent factors representing shared biological programs across samples (**Fig.3E**). We focused on the top 15 latent factors derived using the default parameter settings in the MOFA function (**Fig.3F**). These factors were linked to phenotypic traits (i.e. successful transmission). Among them, three (factor 2, factor 6, and factor 11) significantly differentiated H3-from H1-containing infections (**Fig.3G)**. Notably, only factor 2 showed a strong and consistently association with transmissibility at both 2 and 4 dpi (**Suppl.Fig.3B**), indicating that this biological program most strongly contributes to the pro-transmission phenotype. Genes with the highest positive loadings on factor 2, including *atp2b2, adcy4, grin1, grik4, grm1, hrh1*, and *atp1a3*, were enriched in pathways related to neuroactive ligand-receptor signaling, calcium and cAMP second-messenger regulation, and secretory gland function (**Fig.3H**). These pathways converge on key mechanisms governing mucociliary clearance: calcium–calmodulin and cAMP signaling regulate ciliary beat frequency, while ion transport pathways modulate epithelial fluid secretion and mucus release. The presence of receptors for histamine (*hrh1*), glutamate (*grm1, grik4*), and serotonin (*htr7*) suggests a neuroimmune neuroimmune signaling contributes to epithelial activation during H3-containing infection, consistent with clinical features such as increased nasal secretions during transmissible influenza infection.

To directly link these transcriptional programs to IFNγ-enriched inflammation, we examined correlations between Latent Factor 2 gene signatures and cytokine abundance (**Suppl.Fig.3C**). Latent Factor 2 scores showed a strong and selective association with IFNγ levels in H3-containing infection (**Fig.3I)**. IFNγ-associated genes including *atp2b2*, *tmem163*, *gm7854*, and *vmn2r7* clustered preferentially in H3-infected samples, implicating calcium-dependent ion transport, epithelial secretory activity, and neuroimmune signaling in shaping a mucosal environment optimized for viral expulsion.

Together, these integrated analyses demonstrate that H3-containing infection induces an IFNγ-driven gene network that engages epithelial, neuronal, and electrolyte transport pathways to enhance mucociliary expulsion, establishing a mechanistic link between inflammation, mucus secretion, and increased transmissibility.

### IFNγ regulates shedding and transmission efficiency

The transcriptional and integrative analyses above identified IFNγ as a leading inflammatory signal associated with enhanced shedding during H3-containing infection. To directly test whether IFNγ specifically regulates shedding and transmission, and to distinguish the relative contributions of type-I versus type-II IFNs, we next performed functional studies in mice deficient in IFNγ (IFNγ⁻/⁻) or type-I IFN receptor (IFNAR⁻/⁻).

WT, IFNγ⁻/⁻, and IFNAR⁻/⁻ mice were infected with an H3-containing virus, and monitored for shedding and transmission. IFNAR ⁻/⁻ mice exhibited shedding titers and transmission rates comparable WT controls, whereas IFNγ⁻/⁻ mice showed a significant reduction in both viral shedding and transmission efficiency (93% → 63%, p<0.05) (**Fig.4A**). Importantly, these differences occurred despite comparable index URT viral titers across genotypes **(Fig4A, middle**), indicating that IFNγ specifically regulates viral release than replication.

**Figure 4:**
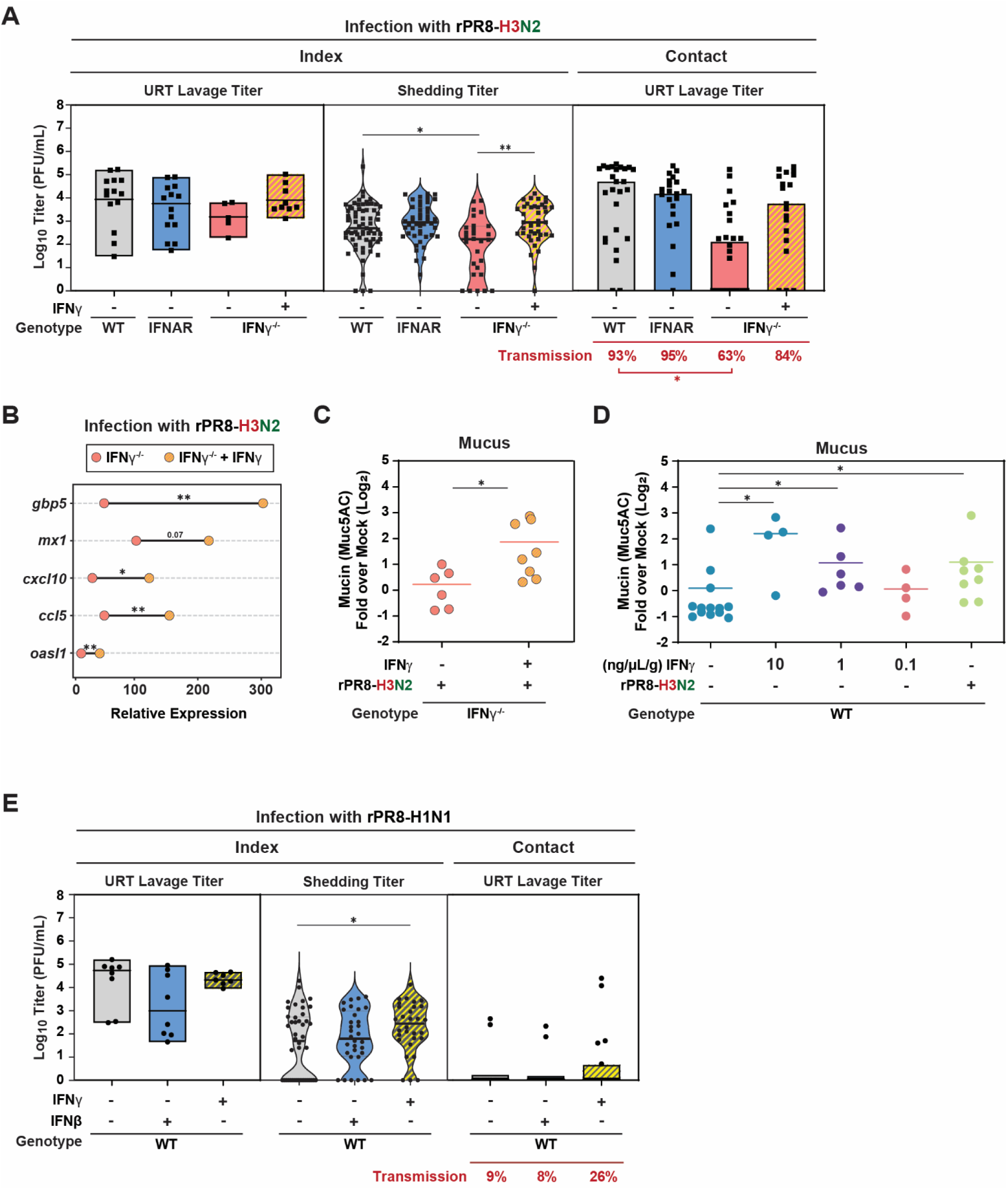
IFNγ regulates viral shedding and transmission efficiency. (**A**) <u>IFNγ supplementation in rPR8-H3N2-infected pups alters shedding dynamics</u>. C57BL/6J (WT) or IFNγ⁻/⁻ index pups were intranasally infected with 300 PFU of rPR8-H3N2 and cohoused with uninfected contacts for 4 days. Index pups received intranasal IFNγ supplementation (1 ng/μL) or vehicle control beginning 4–6 hours post-infection and daily thereafter. Viral titers from day 4 URT lavage (left), cumulative shedding from days 1–4 (middle), and day 4 URT titers from contact pups (right) were quantified by plaque assay. (**B**) <u>IFNγ-dependent transcriptional responses in the URT</u>. Expression of selected antiviral and inflammatory genes in URT cellular fractions was quantified by qRT–PCR at 4 dpi. (**C**) <u>IFNγ restores mucus production in IFNγ⁻/⁻ mice</u>. URT secretions from IFNγ⁻/⁻ pups infected with rPR8-H3N2 and treated intranasally with IFNγ or vehicle control were collected at 2 dpi. URT lavages collected at 2 dpi were dotted on nitrocellulose and probed with anti-Muc5AC antibody. Signal intensity was normalized to positive control, and fold-change over mock is shown. Median values are indicated. (**D**) <u>IFNγ induces mucus secretion in the absence of infection</u>. URT secretions from uninfected pups treated intranasally with escalating doses of IFNγ (10 ng/μL, 1 ng/μL, 0.1 ng/μL) or vehicle, as well as from rPR8-H3N2-infected pups, were collected at 2 dpi. URT lavages collected at 2 dpi were dotted on nitrocellulose and probed with anti-Muc5AC antibody. Signal intensity was normalized to positive control, and fold-change over mock is shown. Median values are indicated. Statistical differences among groups were assessed using the Kruskal–Wallis test. (**E**) <u>IFNγ enhances shedding during rPR8-H1 infection</u>. Index pups infected with 300 PFU of rPR8-H1 received intranasal IFNγ (1 ng/μL/g), IFNβ (1 ng/μL/g) or vehicle beginning 4–6 hpi. Viral titers from day 4 URT lavage (left), cumulative shedding from days 1–4 (middle), and day 4 URT titers from contact pups (right) were quantified by plaque assay. <u>Statistics</u>. Differences between two groups were analyzed using the Mann–Whitney test. Differences among multi-group medians were analyzed by Kruskal–Wallis test. Experiments represent ≥2 independent biological replicates. Significance is denoted as *P < 0.05. Cartoons created with BioRender.com; graphs generated using GraphPad Prism 10.

Reduced shedding in IFNγ⁻/⁻ mice was accompanied by a blunted inflammatory response. Expression of IFNγ-responsive and inflammatory genes, including *gbp5, mx1, cxcl10, ccl5*, and *oasl1*, was significantly diminished relative to IFNγ⁻/⁻ mice supplemented with IFNγ intranasally (**Fig.4B**), resembling those observed during H1-containing infection (**Fig.3D**). Consistent with these transcriptional changes, quantification of mucins in nasal lavages also revealed that IFNγ supplementation in H3-infected IFNγ⁻/⁻ mice significantly increased Muc5AC levels compared with untreated counterparts (**Fig.4C)**. Moreover, IFNγ supplementation induced dose-dependent Muc5AC secretion in the URT, with the intermediate dose (1 µg/mL) achieving levels comparable to those observed during H3-containing infection (**Fig.4D)**. These findings demonstrate a non-redundant role for IFNγ in establishing the pro-inflammatory state associated with efficient shedding, and suggests that IFNγ is sufficient to promote mucosal secretory responses independently of viral replication.

To test whether IFNγ signaling could enhance shedding in the context of a low-transmissibility (H1-containing) virus infection, WT mice were infected with an H1-containing virus and treated with recombinant IFNβ or IFNγ starting 4–6 hours post-infection. IFNβ-treated exhibited shedding and transmission rates comparable to untreated controls, despite comparable index URT viral titers (**Fig.4E**). IFNγ-treated mice displayed significantly increased shedding relative to both IFNβ-treated and untreated groups, demonstrating that IFNγ can elevate virus shedding even during an otherwise poorly transmissible infection.

Collectively, these results identify IFNγ as a key regulator of influenza contagiousness. IFNγ orchestrates a hyperinflammatory, hypersecretory mucosal state in the URT that promotes efficient viral expulsion and enhances transmission, independent of effects on viral replication.

## Discussion

IAV transmission is shaped by a complex interplay between viral factors and host physiology, particularly within the URT where shedding initiates spread. While prior work emphasized viral genetics and receptor tropism as determinants of transmissibility and host range, our findings highlight the underappreciated contribution of host inflammation, specifically IFNγ–driven responses, in defining transmission potential^22^. HTVs elicited a distinct, host-driven mucosal state marked by robust IFNγ signaling, epithelial activation, and mucus hypersecretion. This inflammatory environment facilitated vigorous expulsion of virus-laden secretions without necessarily increasing URT replication, reframing transmission as a consequence of how the host responds to infection rather than solely how efficiently the virus replicates. Consistent with this concept, studies in the ferret model have shown that the magnitude and timing of the host local innate immune responses correlate with the frequency of influenza transmission and symptom burden^5^. Together, these insights support a model in which transmission is not just a viral trait but a host-modulated outcome.

Using the infant mouse model, which recapitulates the high URT susceptibility and efficient contagion observed in children^23–25^, we found that H3N2 strains, induced greater shedding and contact transmission than H1N1, despite comparable URT viral titers. These findings align with human epidemiological data showing H3N2 strains are historically more transmissible than H1N1^1,17–19^. Furthermore, these differences persisted across experiments using isogenic recombinant viruses, implicating viral surface proteins, particularly HA, in tuning host inflammatory responses. Although receptor binding preference can influence host range, altering α2,3-versus α2,6-linked sialic acid interactions had only modest effects on transmission, indicating that post-entry host responses, including the intensity of inflammation, are key determinants of contagiousness (i.e. whether an infected host will shed enough virus to infect others)^5^.

Bulk transcriptomic and cytokine analyses revealed that H3-containing infections consistently engaged IFN-stimulated gene networks, with IFNγ emerging as a central regulator of the pro-transmission mucosal state. IFNγ-deficient mice exhibited markedly reduced shedding and transmission despite comparable URT viral loads, and these deficits were rescued by IFNγ supplementation. Thus, IFNγ promotes a hyperinflammatory, hypersecretory URT environment that increases virus-laden mucus outflow.

Our integrative multi-omics analysis further illuminated the biology linking IFN responses to enhanced transmission. MOFA identified a latent transcriptional factor, Factor 2, that was strongly associated with a transmissible phenotype at both 2 and 4 dpi. In contrast to classical antiviral ISGs, Factor 2 genes mapped predominantly to pathways that regulate mucociliary clearance, epithelial secretion, and neuroimmune communication, including calcium and cAMP second-messenger control (*atp2b2, adcy4*), ion transport (*atp1a3*), and neurotransmitter receptor signaling (*grin1, grik4, hrh1*). Together, these findings suggest that IFNγ reshapes the epithelial secretory environment, and potentially neural-epithelial reflexes, to facilitate viral expulsion. Mechanistically, these pathways converge on features clinically recognizable as “cold symptoms”, rhinorrhea, congestion, increased secretions, phenotypes tightly linked to contagiousness in humans. Thus, the host-protective intent of these responses (i.e., rapid clearance of pathogens from the URT) simultaneously generates the conditions most favorable to onward spread. Hence, our results effectively reframe **transmission** as a by-product of the host’s antiviral defense strategy.

The notion that robust mucosal inflammation promotes microbial transmission, is supported by analogous findings in bacterial systems. Diminished nasal inflammation, via immunomodulation or genetic perturbation, reduces pneumococcal shedding and transmissibility independent of colonization density^26^. This underscores the general principle that mucosal inflammation is a key driver of pathogen exit. In our study, H3-containing infection induced a suite of inflammatory mediators, such as IL-1β, TGF-α, and IFNs, that have well-documented effects on airway goblet cells and mucus production. These inflammatory mediators have been shown to upregulate mucin genes like *muc1, muc5ac*, and *muc5b*^9,27,28^, producing a secretion-rich environment optimized for mechanical pathogen clearance. Whether mucin induction reflects direct epithelial sensing of virus or is secondarily driven by immune cell infiltrates remains unresolved. However, our results demonstrate that epithelial remodeling tightly tracks with contagiousness.

The implications of these findings extend beyond the specific virus-host context examined here. Age-specific differences in airway physiology and immunity likely explain why infants and young children are disproportionately efficient spreaders of influenza^23–25^. The infant URT is inherently prone to robust inflammation and mucus-rich responses that protect the LRT by clearing pathogens from the URT, yet simultaneously increase community transmission. Children shed virus in greater quantities and for longer durations than adults, reflecting less mature immune regulation and limited prior immunity. Our infant mouse model recapitulates these features, supporting the concept that a developmentally primed pro-inflammatory mucosal milieu contributes to the superspreading role of children.

In summary, our study our study demonstrates that influenza transmissibility is not solely dictated by viral replication, but is a dynamic outcome critically shaped by host-driven mucosal responses. Highly transmissible viruses leverage a high-inflammation, IFNγ-skewed environment to promote shedding and spread. We propose that this pro-transmission state evolved as an adaptive host response to rapidly clear infection from the URT, via enhanced mucus and reflexive symptoms like sneezing and coughing, thus protecting the LRT. Ironically, these potent innate defenses simultaneously create optimal conditions for contagion as an unintended consequence. By identifying the immune and epithelial pathways that tune host contagiousness, our findings establish an actionable framework for mitigating spread. Targeting the host’s pro-shedding machinery, rather than replication alone, offers a promising avenue to reduce community spread of respiratory pathogens. We can begin to rethink transmission-focused interventions: not just by attacking the virus, but by artfully dialing down the host factors that viruses hijack for their own propagation.

## Material and Methods

### Mice

C57BL/6J mice, B6.129S7-Ifng^tm1Ts^/J and B6(Cg)-Ifnar1^tm1.2Ees^/J (Jackson Laboratories, ME) were maintained and bred in a conventional animal facility. Pups were housed with their dam for the duration of all experiments, and both male and female pups were used. Animal studies were conducted in accordance with the Guide for the Care and Use of Laboratory Animals^29^ and approved by the Institutional Animal Care and Use Committee of NYU Langone Health (assurance no. A3317-01). All procedures were in compliance with the Biosafety in Microbiological and Biomedical Laboratories. This study was approved by the IACUC under protocol # PROTO202200097.

### Cells and viruses

Madin-Darby canine kidney (MDCK) cells (CCL-34, ATCC) cultured in Dulbecco’s modified Eagle’s medium (Cytiva) with 10% fetal bovine serum (Atlas Biologicals) and 1% penicillin-streptomycin (Gibco). They are screened monthly and confirmed negative for mycoplasma. A/X-31(H3N2) (GenBank: OQ925911-18), A/X-47(H3N2) (NR-3663, BEI), A/Hong Kong/1/1968–2_MA21-2(H3N2) (NR-28634, BEI), A/Puerto Rico/8/1934(H1N1) (NR-3169, BEI), A/California/4/2009(H1N1) (NR-13659, BEI), B/Lee/1940 (NR-3178, BEI). IAV and IBV were propagated in 8–10-day-old embryonated chicken eggs (Charles-River, CT) for 2 days, 37°C and 33°C, respectively. Allantoic fluid was ultra-centrifuged (10,000 rpm, 30 min, 4°C) and stored at −80°C. Virus titers were quantified via standard plaque assay in MDCK cells in the presence of TPCK-trypsin (20233; Thermo-Scientific)

### Cloning

Virus mutants were generated using reverse genetics with plasmids obtained from Adolfo Garcia-Sastre (Icahn School of Medicine at Mount Sinai, NY) and constructed using 0.5ug of eight ambisense plasmids (pDZ-HA, pDZ-NA, pDZ-NP, pDZ-NS1, pDZ-PB1, pDZ-PB2, pDZ-PA, pDZ-NS) in 125uL of OPTI-MEM (GIBCO-Invitrogen). HA plasmids with sialic acid mutations were designed with Twist Biosciences (San Francisco, CA) using a pPol1 promoter. Cationic liposomal reagent Lipofectomine 3000 (3.75μL) and P3000 reagent (8μL) in 125μL of Opti-MEM (Gibco) was added to DNA solution and incubated at RT for 15 minutes. Solution was added to 293T cells in dropwise fashion and incubated at 37C, 5% CO2 for 24 hours. After 24 hours, 1mL of Opti-MEM containing 1ug/mL of TPCK was added to each well. 48hrs post transfection, 100μL of supernatant was collected and inoculated into 7-10 day old embryonated chicken eggs at 37C 5% CO2 for 40hrs. allantoic fluid was harvested, and assay for hemagglutination of 0.5% turkey red blood cells (Charles-River, CT) in V-bottom 96 well plate.

### Virus infection, shedding, and transmission

Pups in a litter (4 to 7 days of age) were infected (index) with a 3μl sterile PBS inoculum without general anesthesia (to avoid direct lung inoculation) by intranasal instillation of 300 PFU of IAV (unless otherwise specified) and returned to the litter at the time of inoculation for the duration of the experiment. Shedding of virus was collected by dipping the nares of each mouse into viral medium (PBS plus 0.3% bovine serum albumin [BSA]) daily, and samples were evaluated via plaque assay. Intralitter transmission was assessed in littermates (contact) at 4-7 dpi. (days 10 to 14 of life) after euthanasia in accordance with institutional guidelines. The pups and dam were euthanized by CO_2_ asphyxiation followed by cardiac puncture, the URT was subjected to a retrograde lavage (flushing of 300μl PBS from the trachea and collecting through the nares), and samples were used for various assays (plaque assay, flow cytometry, Luminex, qRT-PCR, RNA-sequencing). Ratios of index to contact pups ranged from 1:3 to 1:4.

### Compound treatment

Pups were treated intranasally with IFNγ (R&D Systems, Lot #169049915, Minneapolis, MN, USA) or IFNβ (R&D Systems, #8234-MB-010, Minneapolis, MN, USA) at 10-0.1ng/μL diluted in 0.1% PBS/BSA and intranasally daily for the duration indicated.

### Plaque assays

Standard IAV plaque assays were carried out on confluent monolayers of MDCK cells using Oxoid purified agar (Fisher Scientific, OXLP0028B) in the presence of TPCK (tolylsulfonyl phenylalanyl chloromethyl ketone)-treated trypsin (Thermo Scientific). Lung samples were homogenized in PBS, centrifuged at 5000 RPM for 5 minutes, and supernatants were collected and used for plaque assays.

### qRT-PCR

Following a retrograde URT lavage with 300 µl RLT lysis buffer, RNA was isolated (RNeasy kit; Qiagen), and cDNA was generated (high-capacity RT kit; Applied Biosystems) and used for quantitative PCR (SYBR Green PCR master mix; Applied Biosystems). Results were analyzed using the threshold cycle (2^−ΔΔ*CT*^) method by comparison to GAPDH (glyceraldehyde-3-phosphate dehydrogenase) transcription. Values represent the fold change over uninfected. Primers used in (**Supplementary Table 3**).

### URT lavage immunoblots

URT lavages [150 µL, 1:50 in TBS (tris-buffered saline)] were applied to slot blotting apparatus (Minifold-II, Schleicher-Schuell) containing pre-wetted 0.2 µm nitrocellulose membrane (GE10600094; Amersham). After applying vacuum, membrane was blocked with Carbofree buffer (SP-5040-125, Vector-Laboratories) overnight (4°C) followed by incubation with biotinylated Muc5AC (1:10,000) in TBS-T (TBS, 0.1% Tween-20) (2hr, 4°C). Membrane was washed 5×/10 min with TBS-T prior to streptavidin-HRP secondary (1:100,000, 1 h, RT). After five washes/10 min with TBS-T, membrane was developed using SuperSignal West-Femto substrate (34095; Thermo-Scientific) using iBright-Imaging (Invitrogen). Mean gray values were measured with FIJI ImageJ (v.1.53q, NIH). Samples were normalized to a positive control (URT lavage from A/X-31-infected mice) which was included on all blots. Fold change over the average mock values were plotted.

### Immunohistochemistry

Pups were euthanized after 2 dpi. After skin removal, intact heads were fixed in 4% paraformaldehyde (48–72 h, 4°C). Heads were washed in PBS (4°C, 30 min), decalcified (0.125M EDTA, 4°C, gentle shaking, 7 days), dehydrated through graded ethanols, and paraffin embedded (Leica Peloris), and 5-µm, deparaffinized sections were placed onto slides.

For multiplex immunofluorescence staining with Akoya Biosciences® Opal™ reagents, slides were incubated with the first primary antibody and secondary polymer pair and then underwent HRP-mediated tyramide signal amplification with a specific Opal® fluorophore. The primary and secondary antibodies were subsequently removed with a heat retrieval step, leaving the Opal fluorophore covalently linked to the antigen. This sequence was repeated with subsequent primary and secondary antibody pairs and a different Opal fluorophore at each step (see table below for reagent details). Sections were counterstained with spectral DAPI (Akoya Biosciences, FP1490) and mounted with ProLong Gold Antifade (ThermoFisher Scientific, P36935). Semi-automated image acquisition was performed on either a Leica AT2 whole slide bright field scanner at 40X magnification or on an Akoya Vectra Polaris (PhenoImagerHT) multispectral imaging system at 20X magnification using PhenoImagerHT 2.0 software in conjunction with Phenochart 2.0 and InForm 3.0 to generate unmixed whole slide qptiff scans. Image files were uploaded to the NYUGSoM’s OMERO Plus image data management system (Glencoe Software). Figures were prepared with OMERO.figure v4.4 (OME team).

| Antibody | IAV-NP | Ly6g | F480-a |
| --- | --- | --- | --- |
| Source | Southern Biotech,10780-01 | BD, 551459 | CST, 70076S |
| Clone | K5609-V682 | 1A8 (RUO) | D2S9R |
| Retrieval | ER2-20 | ER1-30 | ER2-20 |
| **Blocker** | Rodent Block M | Rodent Block M | Leica |
| **Dilution** | 100 | 150 | 1500 |
| **Time** | 30 mins | 60 mins | 30 mins |
| **Secondary** | Mouse-on-Rodent HRP | Rat HRP polymer 1 step | Rabbit-on-Rodent HRP |
| **Vendor/Cat#** | BioCare, MHRP520G | Biocare, BRR4016H | Biocare/RMR622L |
| **TSA fluor** | 570 | 620 | 480 |
| **Cat #** | Akoya/FP1488001KT | Akoya/FP1495001KT | Akoya/FP1500001KT |
| **Dilution** | 150 | 150 | 200 |
| **Time** | 10 mins | 10 mins | 10 mins |

### Luminex

Cytokines and chemokines in URT lavages were measured using 23-plex array analysis (Bio-Rad, Hercules, CA, USA) and ProcartaPlex Mouse IFN-alpha/IFN-beta assay (Invitrogen, Waltham, MA, USA). All cytokines and chemokines were recorded on a MAGPIX machine (Luminex) and quantitated via comparison to a standard curve. All samples were normalized to those from vehicle treated animals. Heatmaps were generated using SRplot^30^.

### RNA Sequencing and Transcriptomic Analysis

URT lavages were collected from infant mice. Lavage samples were centrifuged to pellet cellular fractions, and total RNA was extracted using the RNeasy Mini Kit (Qiagen, 74106) according to the manufacturer’s protocol, including on-column DNase digestion. Total RNA was quantified using RNA Nano Chips on an Agilent 2100 BioAnalyzer. RNA-seq library preps were prepared using the sparQ rRNA HMR Kit (Quantabio, 95216-096) using the recommended input range of total RNA, followed by 16 cycles of PCR amplification. Final libraries were assessed using High Sensitivity DNA ScreenTape on the Agilent TapeStation 4200. Library concentrations were determined using Quant-iT, and samples were pooled in equimolar ratios. The pooled libraries were sequenced as paired-end 50 bp reads on an Illumina NovaSeq X+ platform using a 10B 100-cycle flow cell, targeting ∼250–300 million reads per sample.

Raw sequencing data (FASTQ files) were uploaded to Illumina BaseSpace and assigned to individual biosamples for downstream processing. Reads were aligned using the RNA-Seq Alignment application (Illumina), which performs quality filtering, adapter trimming, and alignment to the *Mus musculus* reference genome (GRCm38/mm10). This pipeline generated sample-level alignment metrics, read counts per gene, and BAM files for each biosample.

Gene-level differential expression analysis was performed using the RNA-Seq Differential Expression application (Illumina), which imports the normalized count matrix generated during alignment and applies statistical modeling to identify differentially expressed genes between experimental groups. Outputs included normalized expression values, fold changes, statistical significance (p-values and adjusted p-values), and ranked gene lists for downstream pathway analysis. Only genes meeting predefined thresholds for adjusted significance and fold-change were considered differentially expressed. Resulting DE gene tables (log2 fold change, adjusted p-values) were exported and used for all downstream visualization and interpretation. Differentially expressed gene graphs and Gene Ontology (GO) pathway enrichment analyses were plotted using SRplot^30^.

### Multi-Omics Factor Analysis

Multi-Omics Factor Analysis (MOFA) was performed to summarize the high-dimensional transcriptomic data in an unsupervised fashion, identifying latent factors that capture biological variability in transcriptomic data across URT samples. Subsequently, we assessed whether these latent factors are associated with infection phenotype. For factors linked to transmissibility, we further investigated the key genes contributing to these latent factors. The top 100 genes, ranked by absolute loadings for the transmissibility-associated factor, were subjected to KEGG pathway enrichment analysis using clusterProfiler^31^. Gene–pathway relationships were visualized using a Sankey diagram^30^. Additionally, genes contributing to transmissibility-associated factors were integrated with cytokine measurements by calculating their Pearson correlations to assess how cytokine variation aligned with underlying gene programs.

### Flow Cytometry

URT lavages were collected from infected pups and resuspended in FACS buffer (1X PBS, 2% FBS, 1mM EDTA). Cells were stained with Zombie UV™ Fixable Viability Kit (BioLegend, 423107, San Diego, CA) and blocked with TruStain FcX™ PLUS (anti-mouse CD16/32) antibody (BioLegend, 156604, San Diego, CA), for 15 mins in PBS, followed by then stained with fluorescence-conjugated antibodies in FACS buffer for 30 mins on ice. Cells were fixed with 1% PFA before flow cytometric analysis on Cytek Aurora Spectral Analyzer. The following anti-mouse antibodies were used: CD11b-BUV395, clone M1/70 (Biolegend, 563553, San Diego, CA), CD8a-BUV737, clone 53-6.7 (RUO) (BD Biosciences, 612759, San Diego, CA), SiglecF-BV421, clone E50-2440 (BD Biosciences, 565934, San Diego, CA), Ly6g-BV510, clone IA8 (Biolegend, 127633, San Diego, CA), CD45.1-BV605, clone A20 (RUO) (Biolegend, 110738, San Diego, CA), CD45.2-BV605, clone 104 (RUO) (Biolegend, 109841, San Diego, CA), F4/80-BV650, clone BM8 (Biolegend, 123149, San Diego, CA), MHCII-BV711, clone M5/114.15.2 (Biolegend, 107643, San Diego, CA), Ly6C-APCfire810, clone HK1.4 (Biolegend, Cat 128056, San Diego, CA), CD64-PE/Cy7, clone X54-5/7.1 (Biolegend, 139313, San Diego, CA), CD4-AlexaFluor 700, clone GK1.5 (Biolegend, 100430, San Diego, CA), CD31-PE/Dazzle594, clone MEC13.3 (Biolegend, 102525, San Diego, CA), CCR2-FITC clone SA203G11 (Biolegend, 150608, San Diego, CA), EpCam-PerCP-Cy5.5 clone, G8.8 (Biolegend, 118220, San Diego, CA), TCRγδ-APC clone GL3 (Biolegend, 118116, San Diego, CA), NK1.1-Pacific blue, clone PK136 (Biolegend, 108722, San Diego, CA), CD19-BUV661 (BD), CD11c-Spark Blue 550, clone PK136 (Biolegend, 108722, San Diego, CA), CD19-BUV661, clone 1D3 (BD Biosciences, 612971, San Diego, CA), CD11c-Spark Blue 550, clone N418 (BioLegend, 117366, San Diego, CA).

### Statistical analysis

GraphPad Prism (v9.4, CA) was used for all analyses (non-parametric, two-tailed tests at *α* = 0.05).

## Acknowledgements

We thank Ralf Duerr, Christian Marier, Adriana Heguy, and Jeffrey N. Weiser for intellectual contributions to the work; Adolfo Garcia-Sastre for influenza virus rescue plasmids. The Experimental Pathology Research Laboratory (RRID:SCR_017928) is partially supported by the partially supported by the Cancer Center Support Grant P30CA016087 at NYU Langone’s Laura and Isaac Perlmutter Cancer Center. The original multispectral imaging system was awarded through a Shared Instrumentation Grant S10 OD021747. Research was supported by NIH/NIAID K08AI141759 and the American Lung Association Richard Star Emerging Respiratory Pathogen Award to M.B.O.

Conceptualization: S.B. and M.B.O.; Data curation: S.B., H.L.R., L.H., S.Y., C.W.; Formal analysis: S.B., M.B.O., C.W.; Funding acquisition: M.B.O.; Investigation: S.B., H.L.R., C.W., G.C., L.H., S.Y., B.T., M.B.O.; Methodology: S.B., L.H., S.Y., C.W., H.L., M.B.O.; Project administration: M.B.O.; Supervision: M.B.O.; Validation: S.B., M.B.O.; Visualization: S.B., M.B.O., C.W., H.L.; Writing-original draft: S.B., M.B.O.; Writing-review & editing: S.B., H.L.R., L.H., S.Y., C.W., G.C., H.L., B.T., M.B.O.

## Data sharing statement

Data are available upon request.

## SUPPLEMENTARY TABLES AND FIGURES

**Supplementary Table 1:** Infectivity and transmissibility of influenza virus strains in the infant mouse model.

| Virus | Subtype | Index Mice |  |  | Contact Mice |  |  |
| --- | --- | --- | --- | --- | --- | --- | --- |
|  |  | No.<br>infected | URT<br>titer | Shedding<br>titer (D1-4) | No.<br>infected | URT<br>titer | Transmission<br>(%) |
| A/Panama/2007/99 (Pan/99) | H3N2 | 5 | 2.52 | 3.26 | 15/17 | 3.37 | 88% |
| A/Hong Kong/1/1968 (HK/68) | H3N2 | 12 | 4.72 | 3.15 | 17/19 | 4.95 | 89% |
| X-31<br>PR/8+A/Aichi/2/1968 (HA/NA) | H3N2 | 7 | 3.83 | 3.42 | 21/21 | 4.73 | 100% |
| A/Puerto Rico/8/1934 (PR/8) | H1N1 | 4 | 4.63 | 2.49 | 2/10 | 3.22 | 20% |
| A/WSN/1933 (WSN) | H1N1 | 6 | 4.51 | 1.58 | 1/10 | 2.86 | 10% |
| A/California/4/2009 (Cal/09) | H1N1 | 11 | 3.19 | 2.08 | 3/30 | 1.46 | 10% |
| A/Brisbane/59/2007 (Bris/07) | H1N1 | 2 | 4.06 | 2.15 | 1/12 | 0.47 | 8% |
| A/Wyoming/3/2003* | H3N2 | 6 | 0 | 0 | 0/21 | 0 | 0% |
| X101*<br>PR/8+A/Beijing/4/1989 (HA/NA) | H3N2 | 6 | 0 | 0 | 0/9 | 0 | 0% |
| X-187*<br>PR/8+A/Victoria/210/2009 (HA/NA) | H3N2 | 6 | 0 | 0 | 0/6 | 0 | 0% |
| rPR8 | H1N1 | 8 | 4.4 | 2.79 | 3/41 | 2.27 | 7% |
| rPR8-N2 | H1N2 | 10 | 3.98 | 2.62 | 0/21 | 0 | 0% |
| rPR8-H3 | H3N1 | 9 | 4.75 | 3.38 | 21/24 | 4.87 | 87% |
| rPR8-H3N2 | H3N2 | 9 | 4.92 | 3.87 | 20/22 | 4.57 | 90% |
| rPR8-H1 (190E 225D) | H1N1 | 8 | 4.4 | 2.79 | 3/41 | 2.27 | 7% |
| rPR8-H1 (190E 225G) | H1N1 | 6 | 4.10 | 3.08 | 0/22 | 0 | 0% |
| rPR8-H1 (190D 225D) | H1N1 | 4 | 3.97 | 2.79 | 0/12 | 0 | 0% |
| rPR8-H3 (226Q 228G) | H3N1 | 7 | 4.19 | 3.42 | 13/18 | 4.38 | 72% |
| rPR8-H3 (226L 228S) | H3N1 | 9 | 4.75 | 3.38 | 21/24 | 4.87 | 87% |
| rPR8-H3 (30PFU) | H3N1 | 8 | 3.31 | 2.71 | 7/17 | 2.71 | 41% |
| rPR8-H3 (3PFU) | H3N1 | 5/8 | 3.2 | 2.68 | 5/19 | 2.68 | 26% |
| * Virus tested but were unsuccessful at infecting mice. |  |  |  |  |  |  |  |

**Supplementary Figure 1:**
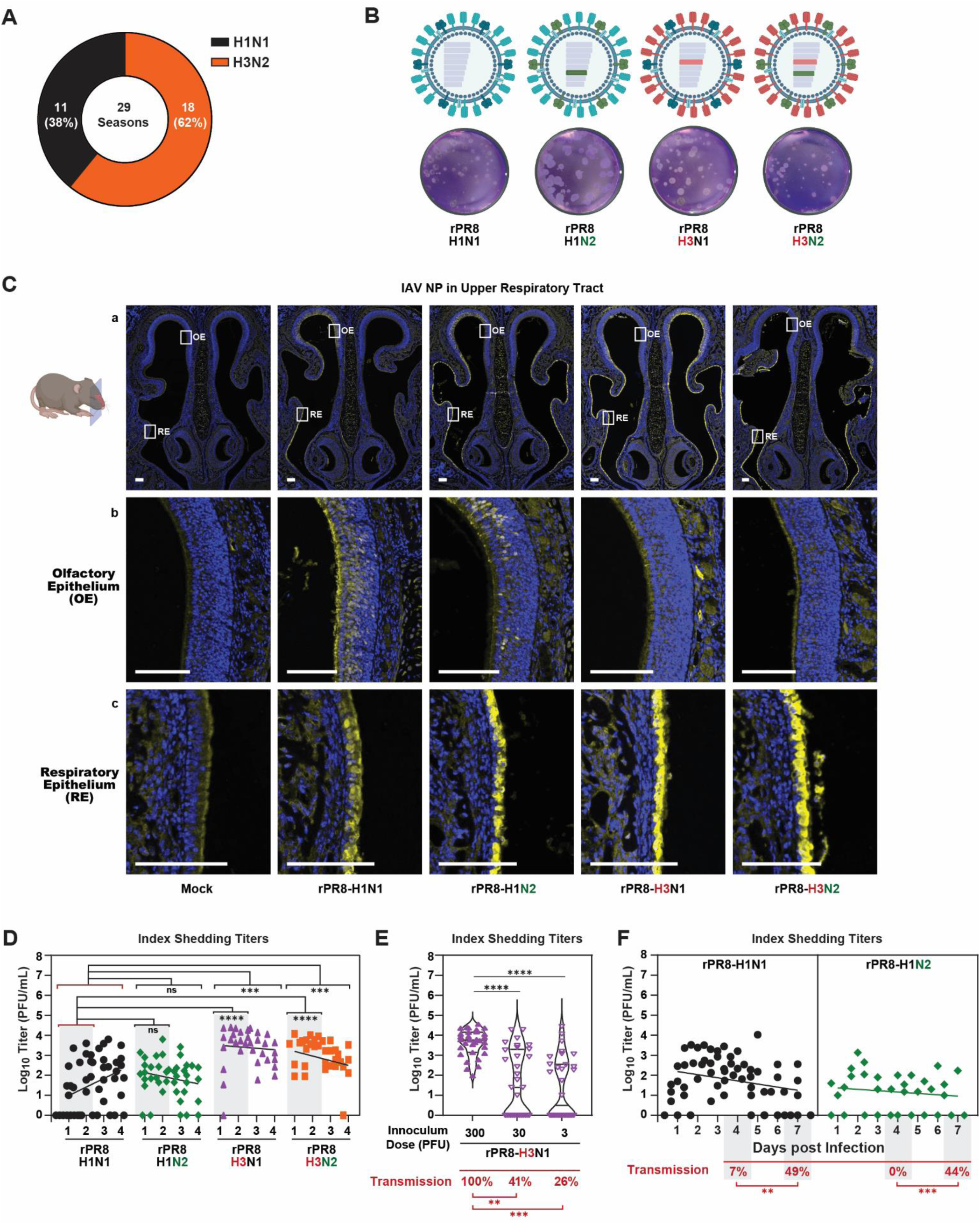
Transmission dynamics of recombinant influenza A viruses with distinct HA and NA combinations. (A) <u>Predominance of H3N2 viruses in human circulation</u>. H3N2 has been the dominant seasonal IAV subtype in 62% of the past 29 influenza seasons in the United States. (B) <u>Schematic of recombinant virus generation</u>. Using reverse genetics, we generated recombinant viruses containing the internal genes of A/PR/8/34 (H1N1) and HA and/or NA segments from A/Hong Kong/1/1968 (H3N2), producing four recombinant viruses: rPR8-H1 (H1N1), rPR8-H3N2 (H3N2), rPR8-H3 (H3N1), and rPR8-H1N2 (H1N2). Viruses were rescued in 293T cells, propagated in 8–10-day-old embryonated chicken eggs, and titrated by plaque assay. (C) <u>Immunohistochemical detection of viral nucleoprotein (NP) in the infant mouse nasopharynx</u>. Infant mice were intranasally infected with 300 PFU of each recombinant virus. At 2 dpi, heads were fixed, paraffin-embedded, sectioned through the nasopharynx, and stained for NP. Expanded views of boxed regions are shown in panels b and c. Scale bars = 100 μm. (D) <u>Daily shedding profiles of recombinant viruses</u>. Shed secretions collected from days 1–4 were quantified by plaque assay. Each point represents an individual mouse. Linear regression curves depict shedding kinetics. (E) <u>Dose-dependent shedding and transmission of rPR8-H3</u>. Index pups were infected with 300, 30, or 3 PFU of rPR8-H3 and cohoused with naïve contacts for 4 days. Daily shedding titers (days 1–4) were determined by plaque assay. Median values are indicated; transmission frequencies are shown in red. (F) <u>Extended transmission analysis of rPR8-H1 and rPR8-H1N2</u>. Index pups infected with 300 PFU of rPR8-H1 or rPR8-H1N2 were cohoused with contacts for 7 days. Shedding was quantified daily (days 1–7). Median values are indicated; transmission rates are shown in red. <u>Statistics</u>. Differences among group medians were analyzed by Kruskal–Wallis test. Data represent ≥2 biological replicates. Significance is denoted as *P < 0.05, **P < 0.01, ***P < 0.001. Cartoons created with BioRender.com; graphs generated using GraphPad Prism 10.

**Supplementary Table 2:** Daily infectivity profiles of index infant mice following recombinant influenza virus infection.

| Virus | Subtype | Index URT Titers |  |  |  |  | Index Shedding Titers |  |  |  |  |
| --- | --- | --- | --- | --- | --- | --- | --- | --- | --- | --- | --- |
|  |  | D1 | D2 | D3 | D4 | D7 | D1 | D2 | D3 | D4 | D7 |
| rPR8 | H1N1 | 2.40 | 5.37 | 4.88 | 4.40 | 4.53 | 1.49 | 2.95 | 2.77 | 2.98 | 2.86 |
| rPR8-N2 | H1N2 | 4.13 | 4.81 | 4.81 | 3.98 | 4.43 | 2.55 | 2.89 | 2.63 | 1.99 | 1.83 |
| rPR8-H3N2 | H3N2 | 5.62 | 6.41 | 5.93 | 4.92 | ---- | 3.33 | 3.73 | 3.12 | 2.86 | ---- |
| rPR8-H3 | H3N1 | 3.87 | 5.03 | 5.41 | 4.75 | ---- | 3.97 | 4.05 | 3.80 | 3.35 | ---- |

**Supplementary Figure 2:**
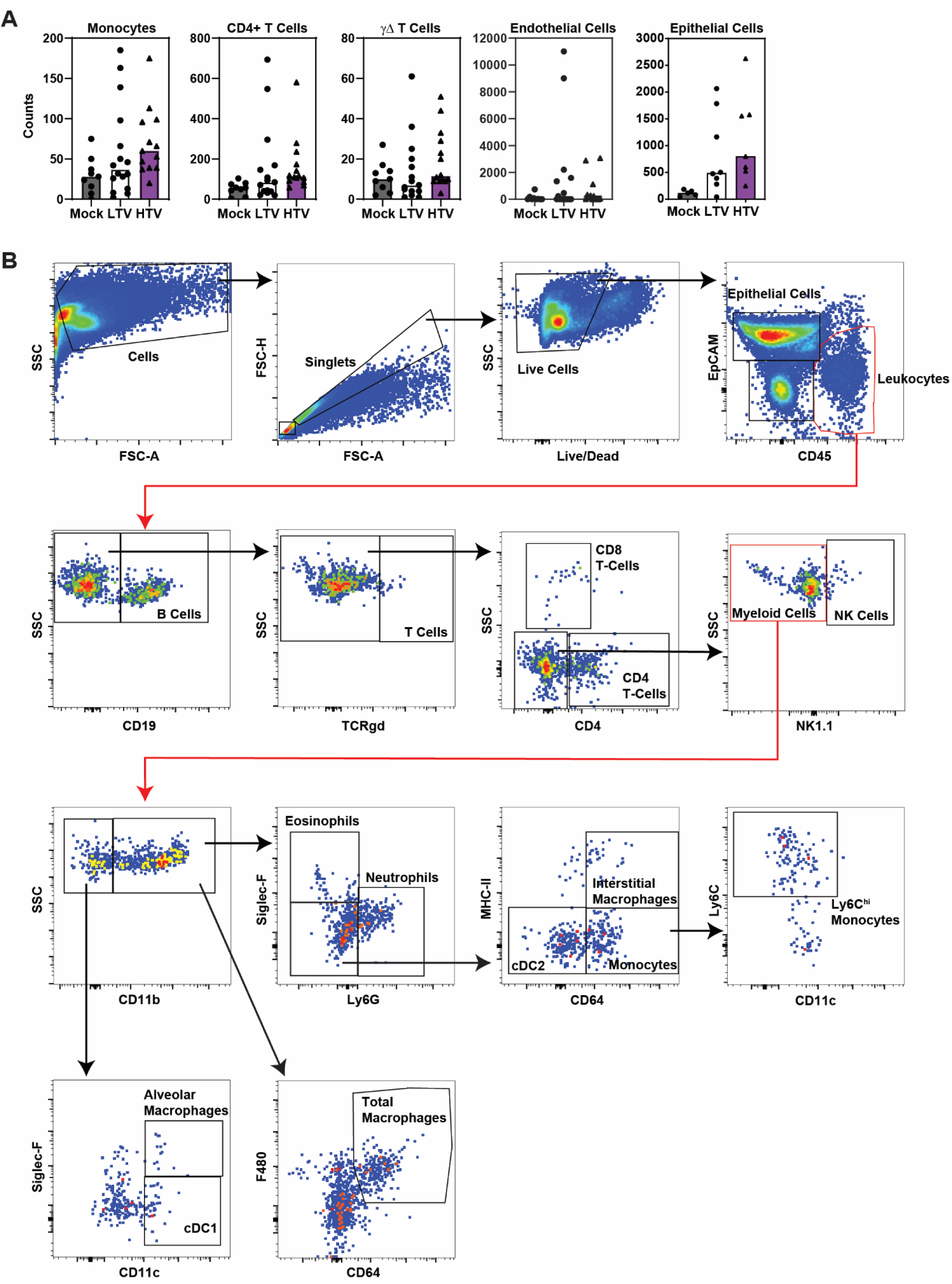
Expanded Flow Cytometry Data. (**A**) <u>Quantification of URT immune subsets</u>. Infant mice were intranasally infected with 300 PFU of rPR8-H3 or rPR8-H1. At 2 dpi, URT lavages were collected and analyzed by flow cytometry. Bar plots show the abundance of monocytes, CD4⁺ T cells, γδ T cells, endothelial cells, and epithelial cells, expressed as a fraction of total live CD45⁺ cells. Data were analyzed using FlowJo and are presented as median values. (**B**) <u>Gating strategy for URT immune populations</u>. Exclusion of debris (FSC × SSC) was done followed by singlets were identified by FSC-H × FSC-A and dead cells (Live/Dead). Epithelial cells were defined as EpCAM⁺CD45⁻. Leukocytes were gated as CD45⁺ cells and further resolved into lymphoid and myeloid subsets. Lymphoid populations included γδ T cells (TCRγδ⁺), CD8⁺ T cells (CD3⁺CD8⁺), CD4⁺ T cells (CD3⁺CD4⁺), and NK cells (NK1.1⁺CD3⁻). Myeloid populations were distinguished as eosinophils (Siglec-F⁺CD11b⁺), neutrophils (Ly6G⁺CD11b⁺), Ly6C^hi^ inflammatory monocytes (Ly6C^hi^CD11b⁺), interstitial macrophages (CD11b⁺MHC-II⁺CD64⁺), cDC2 (CD64^-^MHC-II^-^CD11b⁺), cDC1 (Siglec-F^-^CD11b⁺), alveolar macrophages (AMs; CD11c⁺Siglec-F⁺), and total macrophages (F4/80⁺CD64⁺).

**Supplementary Figure 3:**
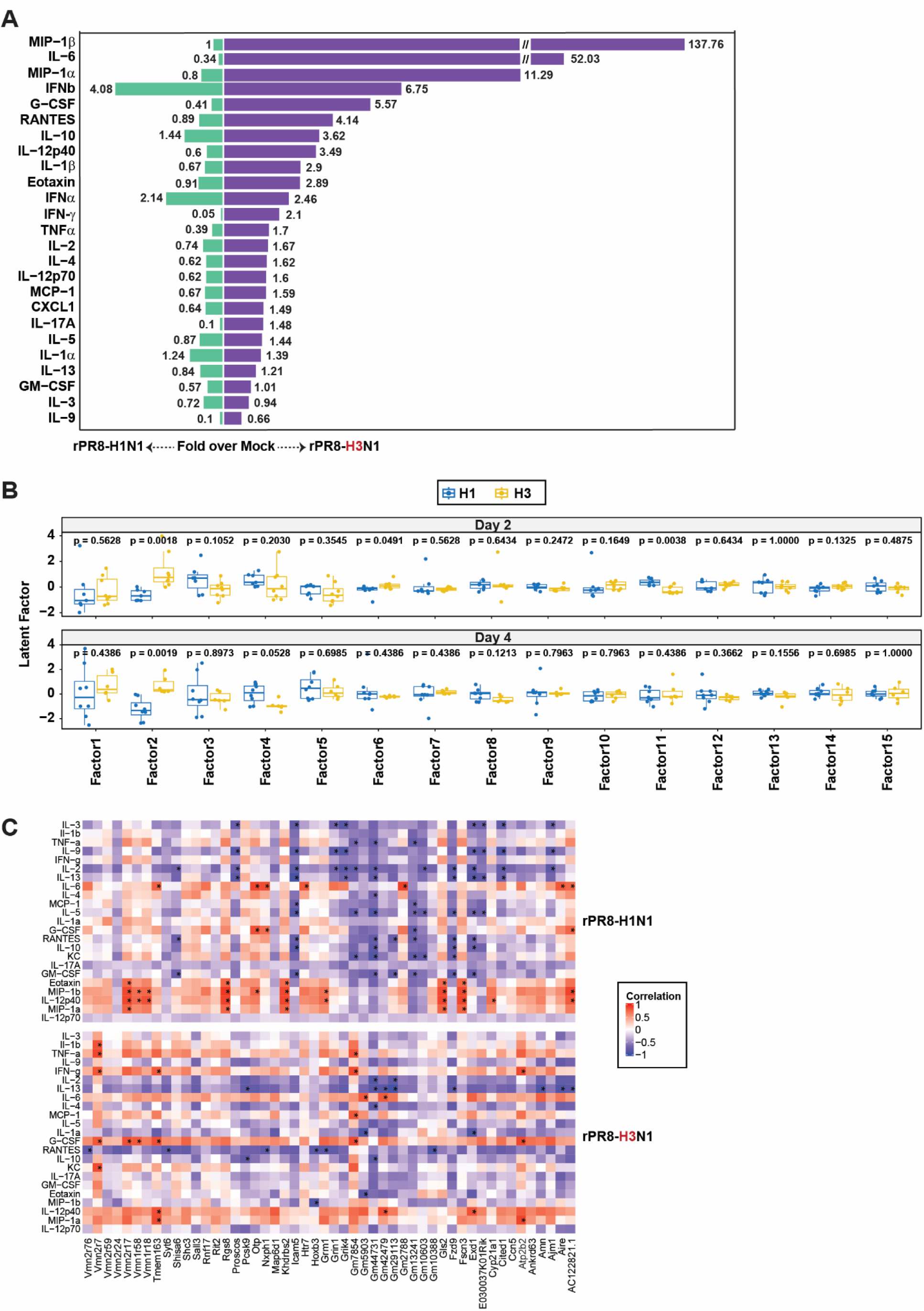
Expanded URT inflammatory and transcriptional profiling. (**A**) <u>Cytokine profiling of rPR8-H3- and rPR8-H1-infected URT secretions</u>. Cytokines shown in Figure 2E were re-graphed to more clearly highlight the magnitude of cytokine induction in URT secretions from rPR8-H3 versus rPR8-H1-infected pups. Data are presented as fold change relative to mock-infected controls. (**B**) <u>MOFA latent factor structure across infection groups</u>. Latent factor structure of the MOFA model. The fifteen latent factors identified by MOFA are displayed according to their distribution in rPR8-H3 and rPR8-H1 infected samples collected at 2 dpi (top) and 4 dpi (bottom). Each plot shows how individual factors segregate between transmission phenotypes, with corresponding p-values. (**C**) <u>Cytokine–gene correlation networks</u>. Heatmap of cytokine–gene correlations. Shown are the correlation coefficients between cytokine abundance and the top 50 genes with the highest positive loadings on Factor 2, the factor most strongly associated with transmissibility. This analysis demonstrates the tight coupling between cytokine responses and rPR8-H3-associated transcriptional programs.

**Supplementary Table 3:**
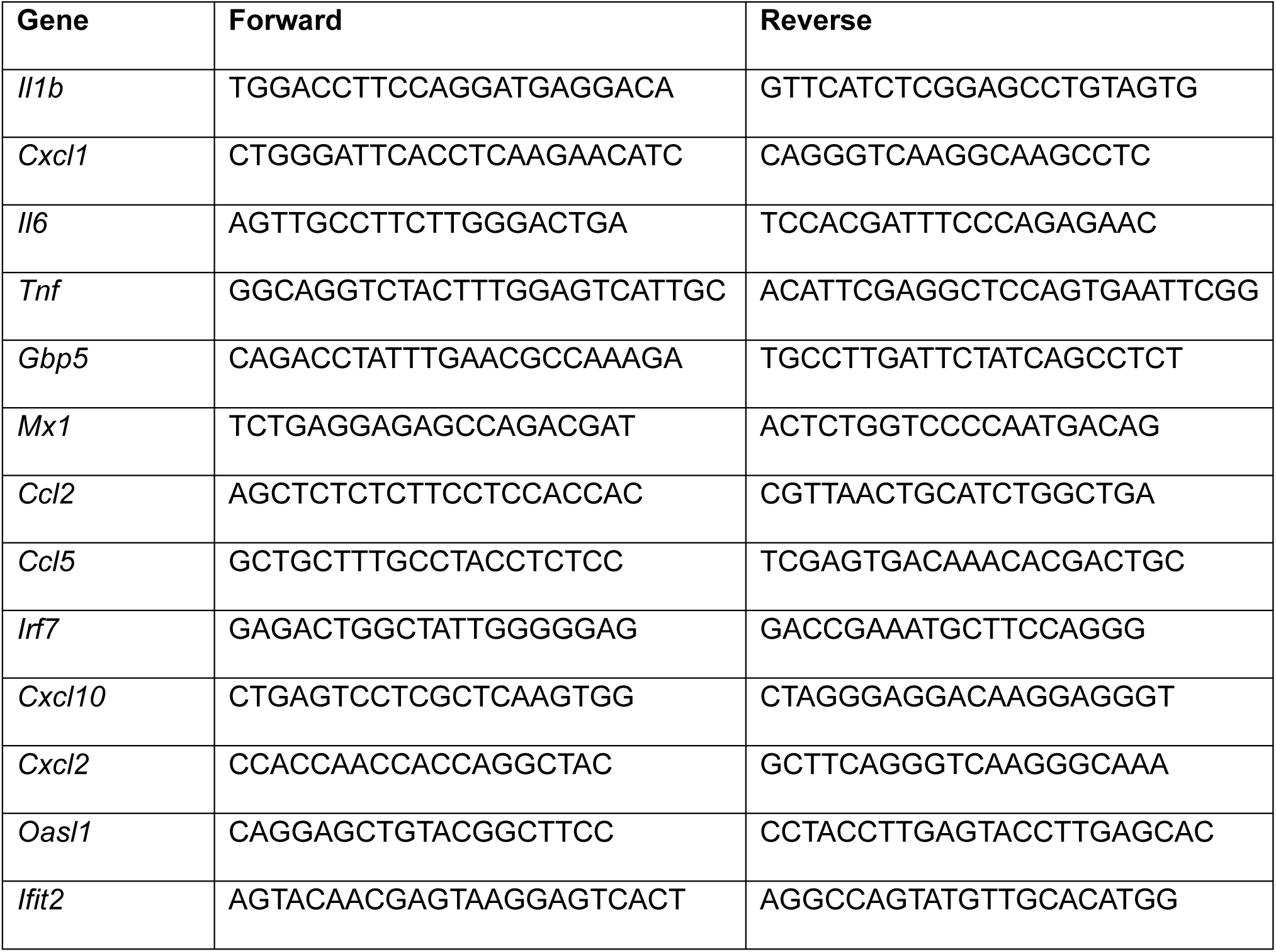
Primers used for SYBR green qPCR.

## Notes

### Competing Interest Statement

The authors have declared no competing interest.

